# Blood clot fracture properties are dependent on red blood cell and fibrin content

**DOI:** 10.1101/2020.10.05.326165

**Authors:** Behrooz Fereidoonnezhad, Anushree Dwivedi, Sarah Johnson, Ray McCarthy, Patrick McGarry

## Abstract

Thrombus fragmentation during endovascular stroke treatment, such as mechanical thrombectomy, leads to downstream emboli, resulting in poor clinical outcomes. Clinical studies suggest that fragmentation risk is dependent on clot composition. This current study presents the first experimental characterization of the fracture properties of blood clots, in addition to the development of a predictive model for blood clot fragmentation. A bespoke experimental test-rig and compact tension specimen fabrication has been developed to measure fracture toughness of thrombus material. Fracture tests are performed on three physiologically relevant clot compositions: a high fibrin 5% H clot, a medium fibrin 20% H clot, a low-fibrin 40% H clot. Fracture toughness is observed to significantly increase with increasing fibrin content, i.e. red blood cell-rich clots are more prone to tear during loading compared to the fibrin-rich clots. Results also reveal that the mechanical behaviour of clot analogues is significantly different in compression and tension. Finite element cohesive zone modelling of clot fracture experiments show that fibrin fibres become highly aligned in the direction perpendicular to crack propagation, providing a significant toughening mechanism. The results presented in this study provide the first characterization of the fracture behaviour of blood clots and are of key importance for development of next-generation thrombectomy devices and clinical strategies.

## 1. Introduction

Acute Ischemic Stroke (AIS), due to embolic occlusion of a cerebral artery, results in over 5.5 million deaths each year [1]. Intra-arterial Mechanical Thrombectomy (MT) is a minimally invasive procedure for treatment of AIS in which the obstructing thrombus is removed using a stent-retriever and/or an aspiration catheter. Based on the recent clinical studies [2,3,12–16,4–11] the proportion of patients experiencing successful procedural revascularization (TICI >=2b) ranged from 76% [17] to 85.4% [18] after all procedures. Although complete revascularization (TICI =3) is achieved in less than 61% of cases [19], and it often takes multiple attempts to remove the complete thrombus. Many investigators attempt to improve interaction between thrombectomy devices and thrombus to increase the successful revascularization rates. Despite improvements from first-generation thrombectomy devices, distal embolism due to clot fragmentation remains a significant challenge for MT procedures. In general, all endovascular MT techniques and devices carry a significant risk of thrombus fragmentation and subsequent distal emboli with associated adverse clinical outcomes [20–23].

Clinical data suggest that thrombus fragmentation and consequent distal emboli are related to thrombus composition. From histological analysis of thrombi retrieved from 85 AIS patients, Kaesmacher et al [20] found a higher fraction of RBC in patients with multiple embolizations resulting from periprocedural thrombus fragmentation. Gengfan et al [24] found that higher thrombus density in nonenhanced computed tomography, associated with higher fraction of RBC, is an independent predictor for secondary embolism. In contrast, Sporns et al concluded that secondary embolism is more probable for clot with lower RBC content [25]. Moreover, per-pass analysis of AIS thrombi show that the composition of thrombus fragments retrieved with each pass of a device during MT are different, further supporting the dependence of clot fragmentation on clot composition [26].

Previous studies have investigated clot hyperelastic and viscoelastic behaviour using compression testing [27–30] and tensile testing [18,31–34]. However, no study to date has investigated the fracture properties of blood clots. The characterisation of fracture toughness of blood clots has the potential to guide the design of improved next-generation MT devices and to inform clinical strategies. Such fracture mechanics investigation for a range of clot compositions may provide insight into the reported clinical link between RBC content and risk of distal emboli generation during MT.

The objective of the current study is to provide the first characterization of composition-dependent fracture behaviour of blood clot. A bespoke experimental procedure for fabrication and fracture testing of compact tension blood clot specimens is developed. Based on histological studies of the composition of clots removed from stroke patients by MT [35–39], tests are performed on three different clot analogue fracture specimen compositions, ranging from high red blood cell (RBC)/low-fibrin clots to low RBC/high-fibrin clots. In addition to experimental fracture testing of blood clots, finite element cohesive zone fracture models are developed to simulate experimental tests and determine fracture strength and fracture toughness as a function of clot composition. Furthermore, this combined experimental-computational fracture mechanics investigation suggests that alignment of the network of fibrin fibres provides a toughening mechanism in blood clots. This provides a mechanistic explanation for the observed dependence of clot fracture toughness on fibrin content.

## 2. Materials and Methods

Developing a reliable test method that is suitable to test the broad range of clot types and compositions can be challenging due to the fragile nature of the clot material. The issue is even more critical when the sample is under loading modes that lead to material stretching, such as fracture testing. Such tests, in contrast to standard compression tests, require gripping of a fragile and slippery material. Another issue for fracture testing of blood clots arises from the high ratio of toughness to strength of clot, resulting in the significant deformation prior to fracture initiation. Finally, consistent fracture testing requires samples that have dimensions far larger than the dimensions of excised AIS clots. Therefore, careful design considerations are required for test specimen fabrication and test-rig design. In this section we outline the development of a novel and bespoke approach to determine the fracture properties of blood clots.

### 2.1 Fracture test

#### Blood clot fabrication

Thrombus material from human sources cannot be readily fabricated into a repeatable specimen geometry for mechanical and fracture testing. Therefore, techniques have been developed to fabricate clinically relevant thrombus analogues in vivo [40]. Clot analogue samples produced from ovine blood have been found to be histologically similar to human clots [27]. Such analogue materials have been shown to exhibit mechanical behaviour that is similar to excised human clots of a similar composition [41], and are commonly used for pre-clinical testing of thrombectomy devices.

The current study uses an established method to fabricate clot analogues using fresh ovine blood [2,42]. Citrated whole blood was used to form platelet-contracted blood clots within 5 hours of blood collection from the donor animals. Three different clot analogues types were fabricated from blood mixtures with 5%, 20% and 40% haematocrit (H). After separation of platelet rich plasma and red blood cell fractions, blood clot mixtures of the desired haematocrit were created. These mixtures were placed in a cylindrical container where the blood mixtures were clotted by adding calcium chloride to reverse the citrate. Finally, the clots were allowed to mature overnight to create flat, disc-shaped clots with a diameter of between 40 and 90mm and thickness of 6 to 9mm. It has previously been established that a decrease in the RBC content of blood clots results in an increase in fibrin content [40].

#### Test method

Test samples were prepared the day after formation by cutting the clots into a rectangular shape with two holes for the purpose of gripping using bespoke stainless-steel punches, (Figure 1a). Firstly, a rectangular sharp punch was used to cut 25×24 mm specimens from the discs of clot (step 1). Another punch was used for making two holes in the rectangle specimen (step 2). A specific design was used for this punch such that it can be fitted on top of the rectangle punch, thereby allowing the precise positioning of the two holes. A bespoke constraining fixture was designed to fit on top of the clot sample, and to allow making notch of the exact size in the desired location in a repeatable way (step 3). A sharp razor blade of 250 µm thickness is used to make a sharp crack in the specimen. Notched specimen with the crack length of *a* as well as the unnotched specimens were prepared (n = 7 for each clot type).

**Figure 1:**
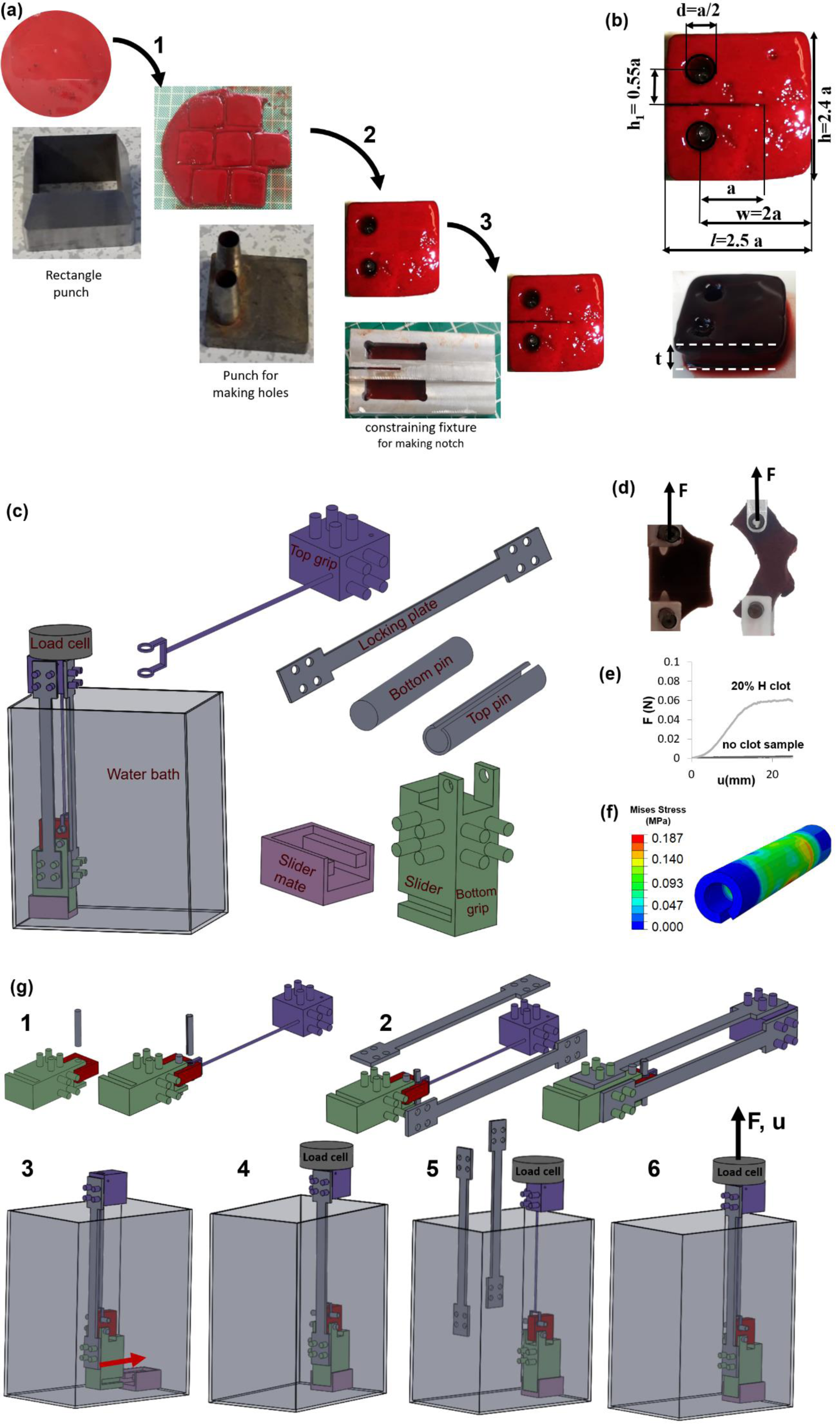
Test method developed for fracture test of blood clot analogues. (a) Notched and un-notched clot specimens were prepared from a large circular clot sample using the bespoke punches. (b) Dimensions of the prepared clot specimens. (c) Designed fracture test rig. (d) Notched and unnotched specimen during the test. (e) The force-displacement curve shows that the hydrostatic force is negligible compared to a representative fracture test result for the developed test rig design. (f) Stress contour for top pin during the fracture test of a 5% H clot for an applied displacement of u=40 mm. (g) Schematic representation of steps for putting the prepared test specimens into the designed test rig.

A bespoke test-rig was developed to perform fracture testing of the prepared samples. A schematic of the designed fracture test rig and each component is shown in Figure 1c. A specific design has been considered for top grip and the pin which connected the clot samples to the grip to minimize the variation of the hydrostatic force and surface tension force during the test. This is a key consideration for this test and any other test which is performed in a fluid medium, especially for the material with relatively low level of force such as clot material and soft biological tissues. The profile area of the top grip was minimised in order to reduce hydrostatic forces that result from submerging the component in a water bath. A hollow bar was designed for the top pin, rather than a solid bar, to minimize the variation of the hydrostatic force during the test. Preliminary validation checks reveal that changes in hydrostatic forces due to upwards movement of the top-gripping fixture are approximately two orders of magnitude lower than the force required to cause fracture (Figure 1e). A finite element rig-design analysis is also performed to ensure that the strength of the top pin provides sufficient mechanical strength during the fracture test. The results as shown in (Figure 1f) reveal that the maximum von-Mises stress in the pin (0.187 MPa) is much lower than the yield strength of the pin (∼210 MPa). A locking fixture is designed to facilitate insertion of the sample into the test rig without inducing pre-test sample damage (Figure 1c). Two locking plates were connected to the top and bottom grips, through the fabricated pins on the grips and prevented any relative movement of grips before test. This design feature is crucial for successful testing of fragile blood clot compositions. Moreover, a slider mechanism is designed for ease of attachment/detachment of the lower grip to the bottom of the water bath without damage to the test specimen. Both bottom grip and its mate component were fabricated from polylactic acid (PLA) using 3D printing.

A schematic of the steps for putting the prepared test specimen into the designed test rig is shown in Figure 1g. Stainless steel bars were placed through the holes of the sample and attached to top and bottom gripping components (step 1). Next, a locking fixture, including two steel locking plates and the pins fabricated on the griping components, was used to prevent the relative movement of the gripping components (step 2). This locking fixture allows confident handling and putting the prepared clot specimen into the test rig without tearing the clot before the test. The prepared samples were then put into a customized experimental setup and fracture tests were performed to determine the mechanical and fracture behaviour of the clot analogue samples. The bottom gripping component slides into an engineered mate component which is glued to the bottom of the water bath (step 3). The top gripping fixture is attached to the mechanical uniaxial Zwick test machine (Zwick Z2.5, Ulm Germany) (step 4). The locking plates are then detached from the gripping components (step 5) and the top grip is moved upwards at a velocity of 10 mm/min and the resultant reaction force is measured throughout the fracture test using a 10 N Zwick load cell (step 6). All tests are carried out in fully hydrated conditions by fully submerging the test specimen and test-rig in a water bath filled with phosphate-buffered saline solution at room temperature (∼20 °C).

#### Size of the test specimen

The choice of specimen dimensions for fracture testing is constrained by the requirement that the initial crack length must be sufficiently large to ensure that crack propagation occurs before the tensile strength is reached in the material [43]. Moreover, the length of the remaining ligament (=W-a) has to be large enough to avoid excessive inelastic deformation in the ligament. There is also a size limit on the thickness of the specimen (t) in relation to the establishment of plane strain or plane stress conditions. More discussion on the specimen thickness is provided in Appendix A. Consequently, large specimens are required for materials with high toughness to strength ratios. This presents a considerable challenge for fracture testing of biological material/soft tissue specimens, where high toughness occurs due to the fibrous nature of the material, yet fabrication of suitably large specimens may not be possible due to the geometric constraints inherent in the natural anatomy of the material. In the current study, we overcome this challenge by using clot analogue materials (rather than excised clots) fabricate large clot analogue samples in order to ensure compliance with the requirements of fracture testing [44]. The so-called critical distance parameter 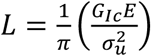, has been used for estimation of the crack length, where *G*_*Ic*_ is fracture toughness, *E* is material stiffness, and *σ*_*u*_ is ultimate strength [45]. The crack length *a* should be in the same order of magnitude as *L* (a larger value of crack length is preferred) to make sure that the crack reaches to the propagation condition before the stress in the specimen reach to the ultimate strength.

Other dimensions of the clot specimen have been chosen based on the length of crack as shown in Figure 1b. It should be noted that the fracture toughness (*G*_*Ic*_) of blood clots has not been reported previously. Therefore we approximate the value of *G*_*Ic*_ for blood clot before the test, based on the value of fracture toughness for soft tissues [45]. We have checked the validity of this initial guess after finishing fracture tests and we found the crack length to be large enough for all three compositions used in this study. Moreover, the clot shows non-linear material behaviour with non-constant stiffness [28,46]. Therefore an approximate values of *E* = 140, 130, 100 *kPa*, which are the values of clot stiffness at 75% strain in unconfined compression test, have been used for 5% H, 20% H, and 40% H clots, respectively. In addition, the value of *σ*_*u*_ = 10.2 *kPa* has been taken from Krasokha et al [47].

Furthermore, we performed a second validity check for the measured fracture toughness. The condition *F*_*max*_*/F*_*Q*_ < 1.1 should be maintained for a valid fracture test, where *F*_*max*_ is the ratio of the maximum load that the specimen was able to sustain during the test [44]. A summary of the validity checks we have done for our fracture test are presented in Table 1 for each clot composition.

**Table 1:** Validity checks for the fracture test of 5% H, 20% H, and 40% H platelet-contracted clot analogues.

|  | 5% H | 20% H | 40% H |
| --- | --- | --- | --- |
| $\frac{F_Q}{F_{max}}$ | 1.07 (<1.10) | 1.02 (<1.10) | 1.00 (<1.10) |
| $L = \frac{1}{\pi} \left( \frac{G_{Ic} E}{\sigma_u^2} \right)$ | 9.37<br>(<a=10mm) | 6.75<br>(<a=10mm) | 2.28<br>(<a=10mm) |

Given the aforementioned size limitations the chosen dimensions for the clot specimen are presented in Table 2 to make sure that the sample is as large as required but also as small as possible. It should be noted that the same dimensions are used for all three compositions, with the exception the specimen thickness dimension, *t*. However, for all cases, the thickness is large enough to ensure plain strain condition behind the crack tip. More details on the effect of specimen thickness are provided in appendix A.

**Table 2:**
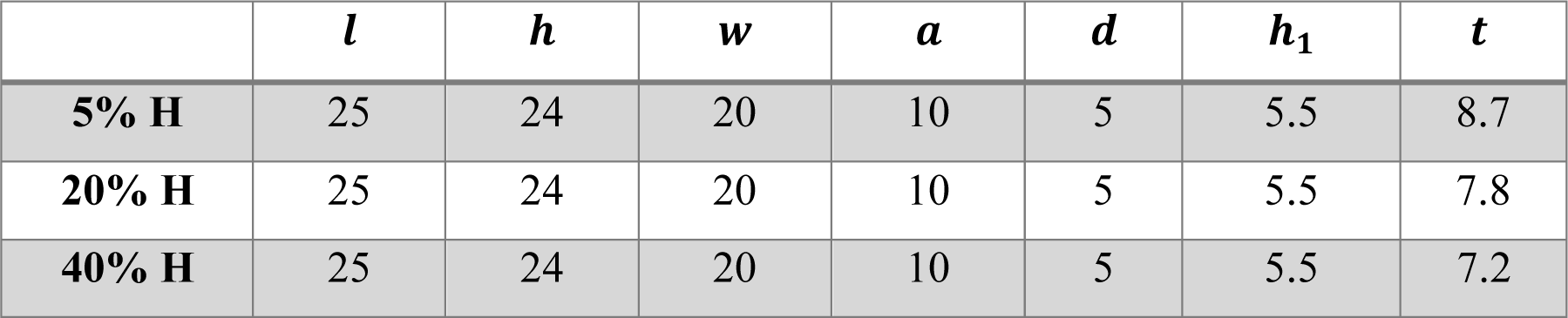
Dimensions of the compact tension clot specimen for fracture test (Figure 1d). All dimensions are in mm.

#### Calculation of the fracture toughness

Fracture toughness which describes the ability of a material to resist the propagation of pre-existing cracks is defined by a parameter *G*_*IC*_ which is the energy needed for crack propagation. The critical strain energy release rate at fracture initiation point, *G*_*IC*_, is calculated form experimental data of fracture test based on the established method for fracture toughness calculation of materials [44]. The method requires determination of the energy derived from integration of the load versus load-point displacement curve in fracture test, while making a correction for the energy which does not contribute to the creation of new surfaces, such as friction losses due to the relative motion of pins and clot. This correction is performed by performing test with an unnotched specimen of the same clot type and subtracting the corresponding energy from that of the notched specimen test (Figure 2). First, the value of a force *F*_*Q*_ is found using the 5% secant method as demonstrated in Figure 2a. The area under force-displacement curve up to the point of *F*_*Q*_ is then calculated which is the energy *U*_1_. Corrected energy *U* is then calculated as *U* = *U*_1_ *− U*_2_, where *U*_2_ is the area under force-displacement curve of the unnotched specimen test up to the point with force *F*_*Q*_. The value of *G*_*IC*_ is then calculated from the corrected energy, *U*, as follows:

**Figure 2:**
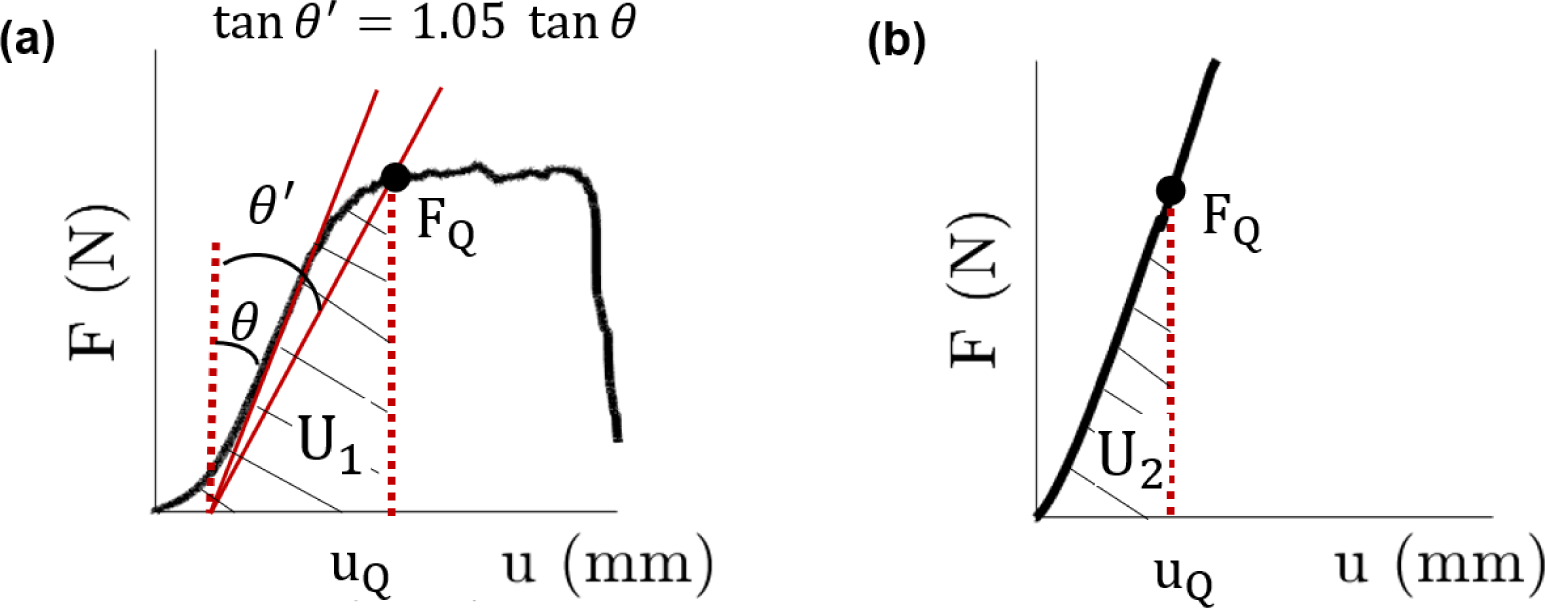
Method for fracture toughness calculation of blood clot. (a) A representative force-displacement curve for fracture test and (b) un-notched compact tension test of blood clot are shown, indicating the work until crack propagation U_2_ and the corresponding energy in unnotched test U_2_ as well as the force F_Q_ and load-point displacement u_Q_ at the start of crack propagation which are used to calculate the fracture toughness (tearing resistance).

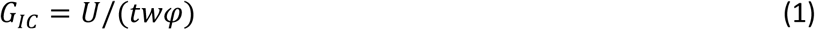

where the energy-calibration factor *φ* is a geometry function depends on the crack length to width ratio (a/w). A value of *φ* = 0.199 is used for a/w=0.5 in this study which is obtained from [48].

### 2.2 Compression test

In order to further characterise the non-fracture biomechanical behaviour of the clot analogue materials, unconfined compression testing is performed using a protocol recently developed by Johnson et al. [27]. In summary, test specimens were prepared by cutting the clots into cylindrical-shaped samples with an approximate diameter of 10 mm, and an approximate height of 5 mm. The testing was performed using a Zwick uniaxial tensile machine (Zwick Z2.5, Ulm Germany) with a customised aluminium compression platen. The clot specimens were loaded to a compressive nominal strain of 80% at a constant axial strain-rate magnitude of 10% per second.

### 2.3 Computational modelling

#### Constitutive modelling and material parameters identification

In order to obtain detailed insights from the fracture tests described above, finite element simulations are performed in which crack initiation and propagation are predicted using a cohesive zone model. The clot material is modelled as an anisotropic hyperplastic fibrous soft tissue using the a recently proposed formulation [49] This formulation has been shown to accurately predict the isochoric and volumetric behaviour of blood clots over a wide range of clot compositions while facilitating control of unphysical auxetic behaviour [28]. The isochoric strain energy density function is given as

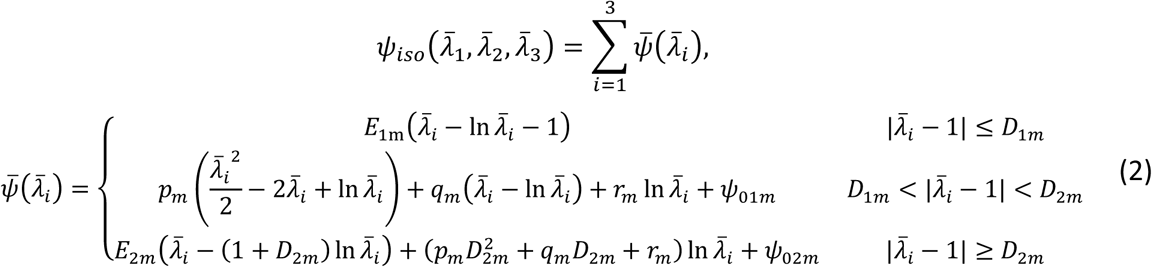

where *D*_1*m*_, *D*_2*m*_, *E*_1*m*_, and *E*_2*m*_ are material parameters, 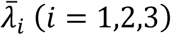 are the isochoric principal stretches, *J* =*λ*_1_*λ*_2_*λ*_3_ is the jacobian, and *ψ*_01*m*_ and *ψ*_02*m*_ are two constants which ensure the continuity of strain energy. Moreover *p*_*m*_, *q*_*m*_, and *r*_*m*_ are not independent parameters; in order to maintain C^0^ and C^1^ continuity the following relations must be enforced:

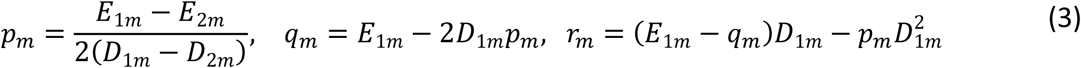

Moreover, the volumetric strain energy density function is given as:

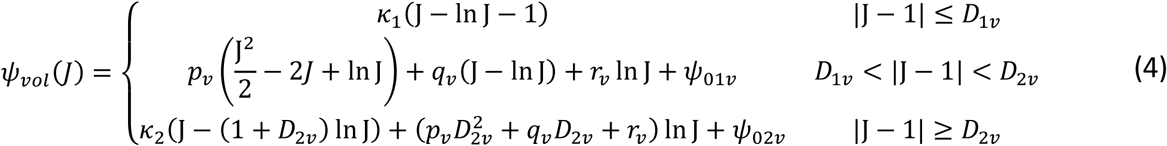

in which *k*_1_ and *k*_2_ are the initial small-strain and large-strain bulk modulus, respectively, the parameters *D*_1*v*_ and *D*_2*v*_ control the transition volumetric strains, and *p* _*v*_, *q* _*v*_, and *r* _*v*_ are obtained in a similar manner as equation (2) by using the corresponding volumetric parameters. For further discussion on the volumetric strain energy density function the reader is referred to the recent paper by Moerman et al. [50].

The anisotropic strain energy density function corresponding to the contribution of fibrin fibres in blood clot is give as:

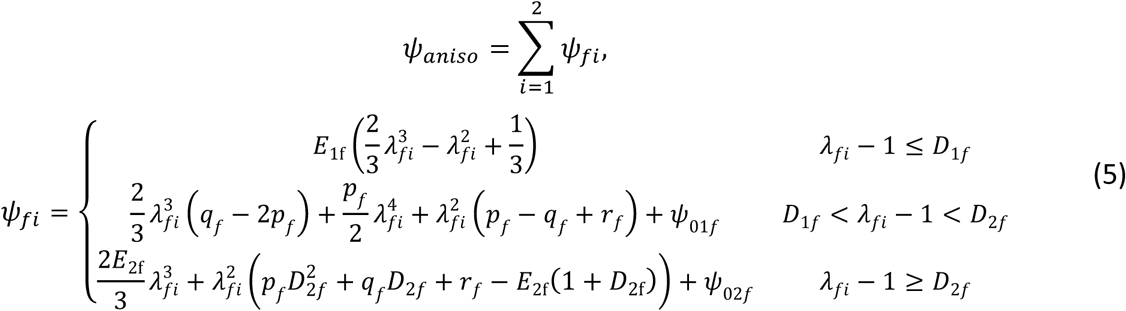

where, and *p*_*f*_, *q*_*f*_, and *r*_*f*_ are obtained in a similar manner as equation (2) by using the corresponding volumetric parameters. The total strain energy density of the clot material is given as 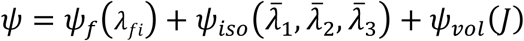. Stress-strain relationships are readily derived from equations (3)-(6) above, as described by Fereidoonnezhad and McGarry [49].

The constitutive parameters for each clot composition are obtained using an inverse finite element parameter identification scheme and least-square optimization algorithm based on the fracture and compression test data. These calibrated material parameters are used for all computational simulations in this study.

#### J-Integral analysis

In 1968, Rice introduced an energetically motivated generalized crack driving force referred to as the J-integral [51]. The crack will grow upon reaching the J-integral to a critical value J_C_. Here, J-integrals for each of the three tested clot compositions are calculated by using the *virtual crack extension method* in Abaqus (2018, Dassault Systémes Simulia Corp.). The computed value of J-integral during the fracture test is then compared to the critical value of the strain energy release rate (G_IC_=J_c_) obtained from the fracture experiment. This provide a good basis for the assessment of the composition-specific fragmentation risk of the clot during clinical procedures such as mechanical thrombectomy.

#### Modelling of crack propagation

The cohesive zone fracture model developed by Fitzgibbon and McGarry [52,53], is used to simulate the crack propagation in the fracture test. In this CZM formulation the magnitude of the interface traction is given as

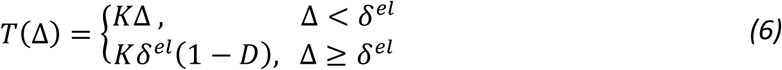

where *K* is the interface stiffness, Δ is the magnitude of the interface displacement vector, and *δ*^*el*^ is the maximum separation prior to the initiation of damage. Interface damage, *D*, increases monotonically from 0, at the onset of damage, to 1, at the point of ultimate failure, such that:

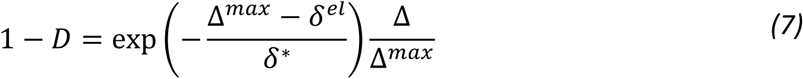

Δ^*max*^ is the maximum displacement; *δ*^∗^ governs the rate at which damage increases with increasing interface displacement. Moreover, the CZM fracture energy *U*_*CZM*_ is introduced as the area under traction-separation curve in the CZM model [53].

The finite element analysis has

.1been performed by using the C3D8 elements in Abaqus/standard. A mesh convergence study was carried out based on the value of force at fracture initiation point and a mesh with 36084 elements was found sufficiently accurate and was used for the calibration of interface strength (*T*) and CZM fracture energy (*U*_*CZM*_). The pins were modelled as rigid surfaces and a frictionless contact between the pins and the clot specimen has been considered.

## 3. Results

### 3.1 Fracture and compression test results

The results of fracture and compression tests for 5%H, 20%H, and 40%H platelet-contracted clot analogues are presented in Figure 3. In general, it is observed that the clot with higher fibrin content are stiffer than the clot with lower fibrin content, both in compression and compact tension test of un-notched specimens. The force-displacement curves of the compact tension fracture test (Figure 3c) exhibit an initial non-linear stiffening followed by stable crack propagation with an approximately constant force. The steady-state force is significantly higher with increasing fibrin content (decreasing haematocrit). The 5% H clot has the greatest peak force at steady-state crack propagation zone, followed by the 20% H clot, with the 40% H clot having the lowest peak force in the crack propagation zone (Figure 3h). The 40% H clot has the lowest initial stiffness, compared to the 20% H and 5% H clots (Figure 3e,f). It is also observed that the crack propagation starts at a higher value of applied deformation (u) for the fibrin-rich clots compared to the RBC-rich clots (Figure 3g). The applied displacement at the final fracture point (*u*_*ff*_) is also higher for 5% H clots compared to the 20% H and 40% H clots. However, the difference in *u*_*ff*_ for 20% H and 40% H clots is not statistically significant (Figure 3i), whereas a statistically significant difference is observed between all clot compositions in terms of the force at fracture initiation ((*F*/*t*)_*fi*_).

**Figure 3:**
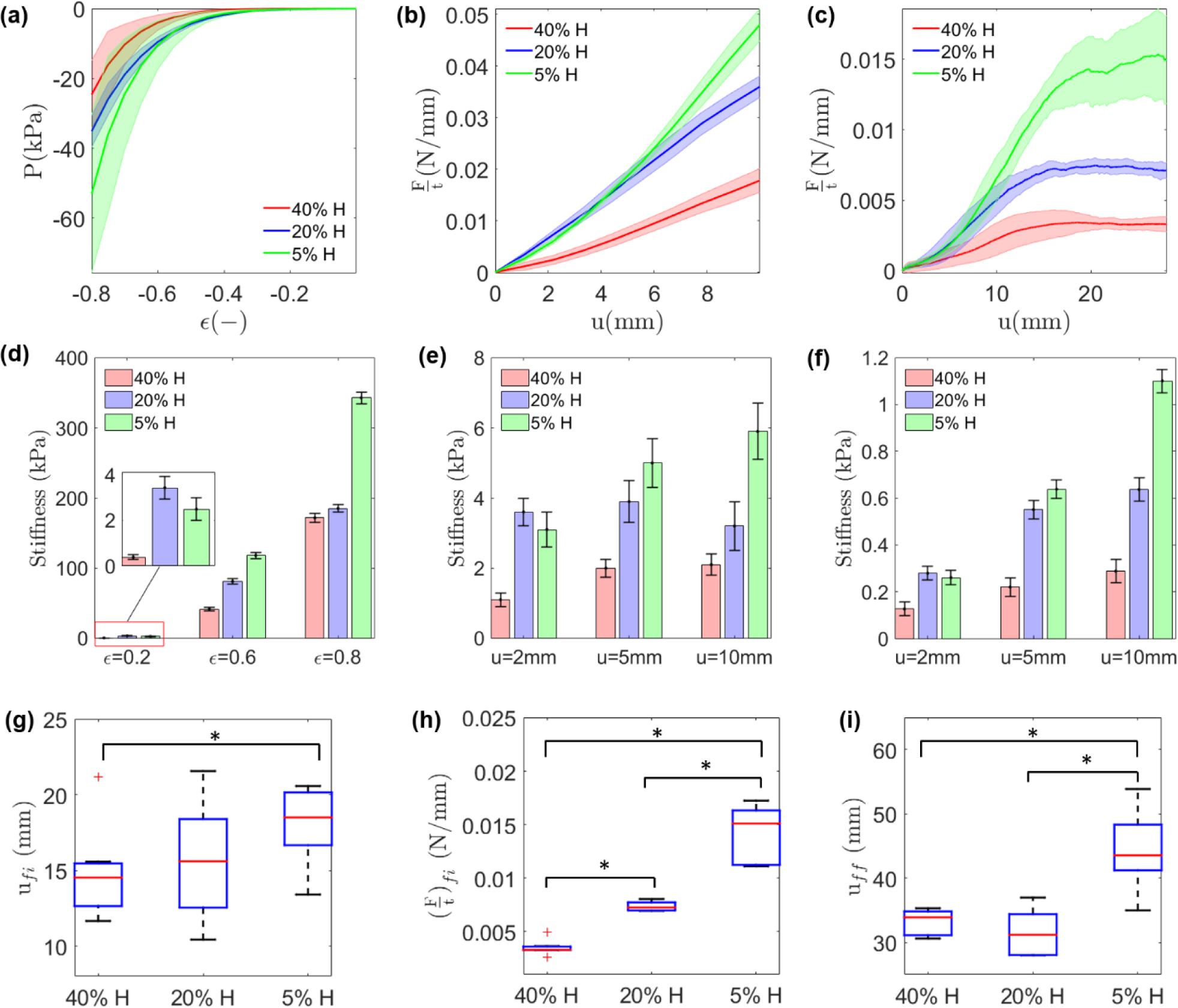
Mechanical and fracture behaviour of clot are significantly dependant on clot composition. Compression, compact tension un-notched test, and compact tension fracture test results for 40% H, 20% H, and 5% H clot analogues. (a) Nominal stress (P) vs. nominal strain (ϵ) for unconfined compression tests; (b) Force per thickness of the specimen (F/t) vs. applied displacement (u) in compact tension un-notched tests. (c) Force per thickness of the specimen (F/t) vs. the load-point displacement (u) in compact tension fracture test; (d) Tangent stiffness at 20%, 60%, and 80% strain in compression test; (e) Tangent stiffness at u=2, 5, 10 mm in compact tension un-notched test; (f) Tangent stiffness at u=2, 5, 10 mm in fracture test; Composition-specific difference in: (g) applied displacement at initiation of fracture (u_fi_), (h) The stable crack propagation force per thickness ((F/t)_fi_), (i) applied displacement at final fracture point (u_ff_). Results are presented as boxplots, where boxes represent upper and lower quartiles and lines inside the boxes define the median, while + represent outliers, and whiskers 10–90 percentiles. Significant differences are indicated for p < 0.05 by * (ANOVA test).

The critical strain energy release rates at crack growth initiation point *G*_*IC*_ for each clot composition are calculated from fracture test results (Figure 3b,c). Figure 4 demonstrates a significant decrease in critical strain energy release rate (fracture toughness) of clot analogous with increasing haematocrit (RBC content); i.e. an increase in fracture toughness with increasing fibrin content. This suggests that fibrin content is a key determinant of the fracture resistance of blood clots.

**Figure 4:**
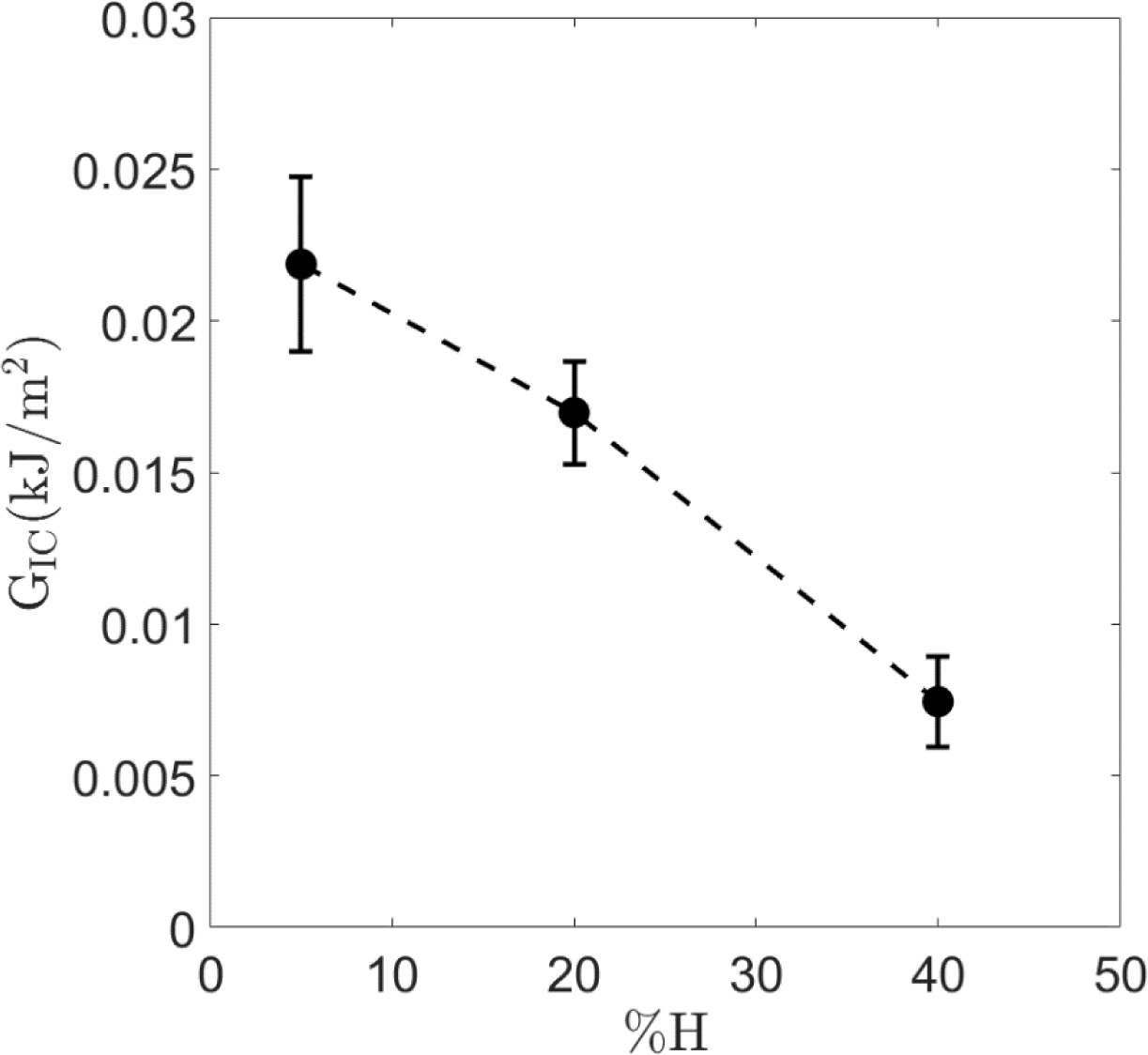
Dependence of fracture resistance on clot composition. The critical strain energy release rate G_IC_ (fracture toughness) at onset of crack propagation for 40% H, 20% H, and 5% H platelet-contracted clot analogues.

### 3.2 Computational results

#### Constitutive parameters identification

The composition-specific constitutive parameters are obtained from compression and compact tension test data and the best-fit parameters are presented in Table 3. The ability of the calibrated constitutive law to predict the mechanical behaviour of different clot types is also demonstrated in Figure 5 where a good agreement between experimental data and model predictions is observed for all clot types. Importantly, if the clot is modelled as an isotropic hyperelastic material a reasonable fit can be obtained for unconfined compression experiments but not for compact tension experiments. However, when an anisotropic component (equation (5)) is incorporated to represent the fibrin network, a reasonable fit can be computed for both unconfined compression experiments and compact tension experiments. This highlights the significant contribution of fibrin alignment and stretching when blood clots are subjected to tensile loading. The anisotropic fibrin component is included in all subsequent simulations in this paper. Furthermore, in appendix B, we have shown that an Ogden hyperelastic model with asymmetric behaviour is not able to capture the experimental results of the unconfined compression test and compact tension test at the same time.

**Table 3:** Best-fit hyperelastic parameters of the constitutive model for 5% H, 20% H and 40% H platelet-contracted clot analogues

|  | 40% H | 20% H | 5% H |
| --- | --- | --- | --- |
| *Matrix parameters* |  |  |  |
| **D1m** | 0.55 | 0.4 | 0.4 |
| **D2m** | 0.65 | 0.55 | 0.55 |
| **E1m (kPa)** | 0.02 | 0.04 | 0.05 |
| **E2m (kPa)** | 8 | 8 | 20 |
| *Fibre parameters* |  |  |  |
| **D1f** | 0.005 | 0.001 | 0.001 |
| **D2f** | 0.01 | 0.002 | 0.002 |
| **E1f (kPa)** | 3 | 32 | 42 |
| **E2f (kPa)** | 15 | 40 | 45 |
| *Volumetric parameters* |  |  |  |
| **D1v** | 0.014 | 0.015 | 0.015 |
| **D2v** | 0.022 | 0.025 | 0.025 |
| **k1 (kPa)** | 2 | 4 | 4 |
| **k2 (kPa)** | 6 | 6 | 6 |

**Figure 5:**
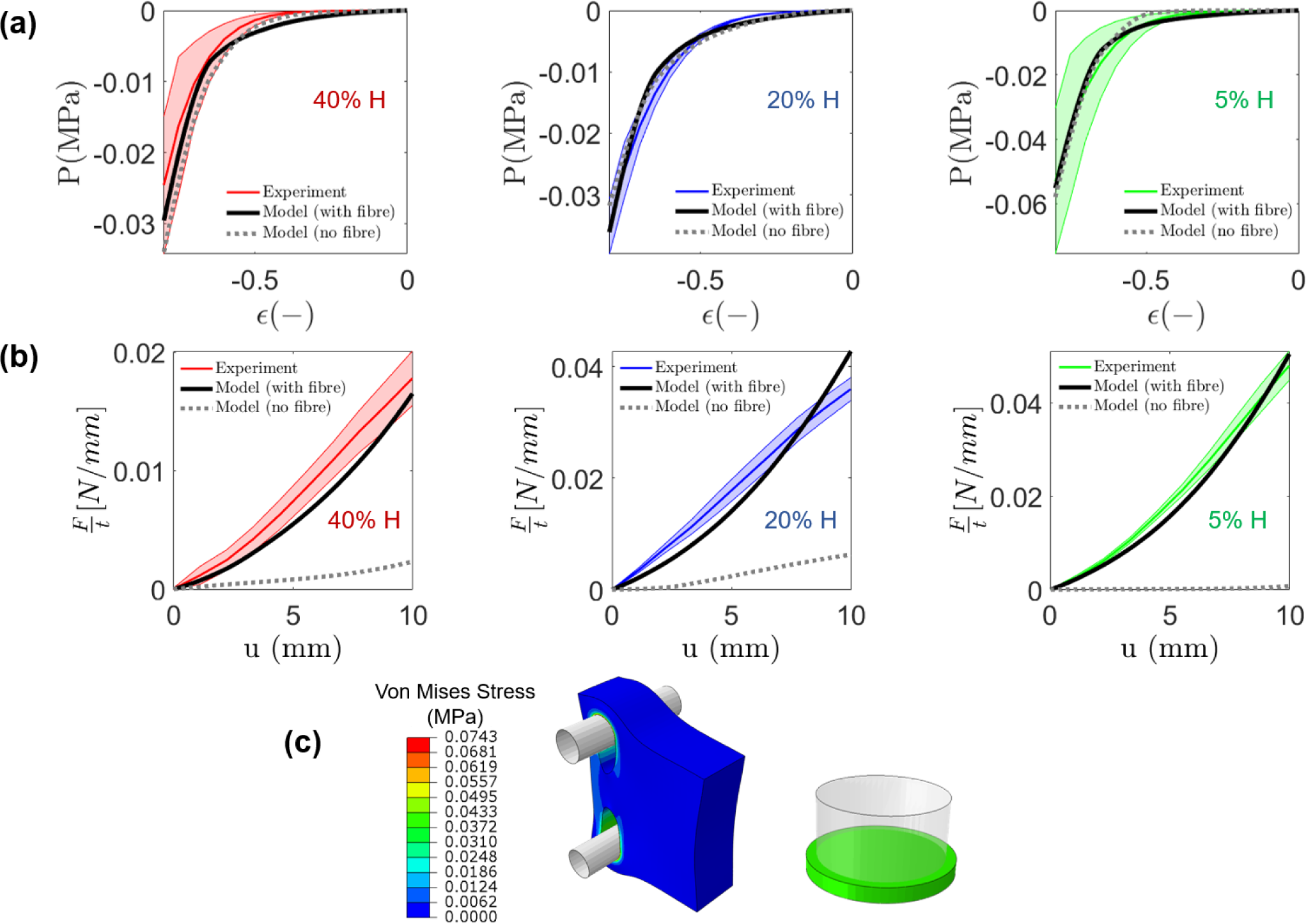
Computational simulation of unconfined compression experiments and compact tension experiments. (a) unconfined compression test; (b) unnotched compact tension test are compared with the experimental data for 5%, 20%, and 40% H platelet-contracted clot analogues; (c) Contour plot of the von-mises stress distribution in a 20% H clot analogue for an unnotched compact tension test simulation at an applied displacement of u=10 mm, and for an unconfined compression test at 80% compression.

#### Role of fibrin fibre alignment

Computed J-integrals are shown as a function of applied displacement (Figure 6a). The 5% H clot has the greatest fracture initiation displacement (u_fi_), followed by the 20% H clot, with the 40% H clot having the lowest fracture initiation displacement.

**Figure 6:**
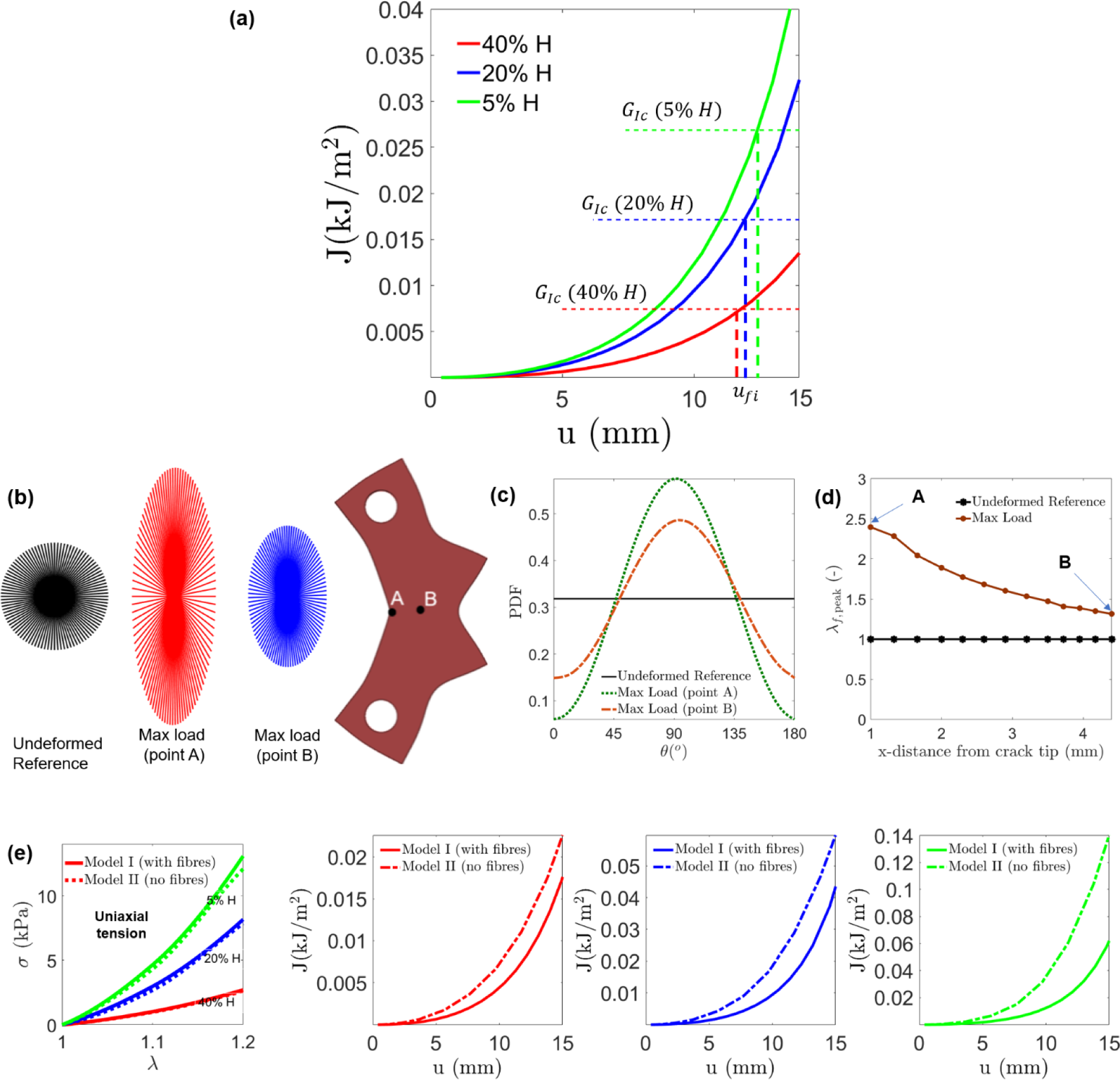
Role of fibrin fibres on the tear resistance of clot. (a) Composition-specific J-Integral calculations for fracture path in the crack direction as a function of the applied displacement (u) in a fracture test. (b) The evolution of fibre alignment behind the crack-tip is shown for a 5% H clot at a point near the crack tip (point A) and a point far from the crack tip (point B). At point A the fibres are highly aligned in the direction perpendicular to the crack propagation direction at maximum load (u=40 mm). (c) Degree of fibre alignment in clot is quantified by the probability density function (PDF) of the fibres as a function of the angle w.r.t the crack direction (θ), for reference (undeformed) state and for the maximum load state and points A and B. (d) The corresponding peak fibre stretch at the notch along the direction of crack (x-direction) is computed. (e) The key role of fibrin fibres in fracture resistance of blood clot is demonstrated by using an anisotropic model which incorporating the influence of fibrin fibres (Model I) and a isotropic model without the fibrin fibres (Model II). Both models have the same uniaxial behaviour.

This computational prediction is in agreement with the experimental observations in Figure 3g. More detailed analysis of the computational simulations reveals fibre alignment perpendicular to the crack direction ahead of the crack (Figure 6b, c). Moreover, the peak value of the fibre stretch (*λ*_*f*,peak_) is higher behind the crack-tip compared to points farther from the crack (Figure 6d). The results in Figure 6 can explain the toughness enhancement in blood clots. It is noted that the fibres are isotropically distributed at the undeformed reference configuration, and they become aligned as the material deforms and stretches in front of the crack tip.

For further investigation on the contribution of fibrin fibres into the fracture resistance of blood clot, we consider two different models: an anisotropic constitutive model which incorporate the contribution of fibrin fibres (model I) and a non-fibrous isotropic model (model II). The hyperelastic parameters of Table 3 are used for model I and the parameters of model II are identified such that both models have the same stress-strain behaviour in uniaxial tension (Figure 6e). The results as demonstrated in Figure 6e reveal the key role of fibrin fibres in fracture resistance of blood clots. Notably, much more deformation is required to reach G_IC_ when the contribution of the fibres are considered (mode I), compared to the model II which does not incorporate the contribution of the fibres. Also, the difference between model I and model II are higher for the 5% H clot with high amount of fibrin compared to the 40% H clot with lower fibrin content.

#### Modelling of crack propagation

FE simulations of the fracture test with notched specimen are performed by using the calibrated constitutive law and CZM model. The results as shown in Figure 7 reveal a good agreement between computational predictions and experimental measurements, in terms of the initial hyperelastic deformation, and in terms of crack initiation and propagation. The interface strength (*T*) of 480 kPa, 250 kPa, and 100 kPa and the fracture energy (*U*_*CZM*_) of 1.152 kN/mm, 0.3645 kN/mm, and 0.045 kN/mm are found for 5% H, 20% H, and 40% H clots, respectively. Similar to the fracture toughness (G_IC_), both interface strength and CZM fracture energy increase by increasing the fibrin content in clot.

**Figure 7:**
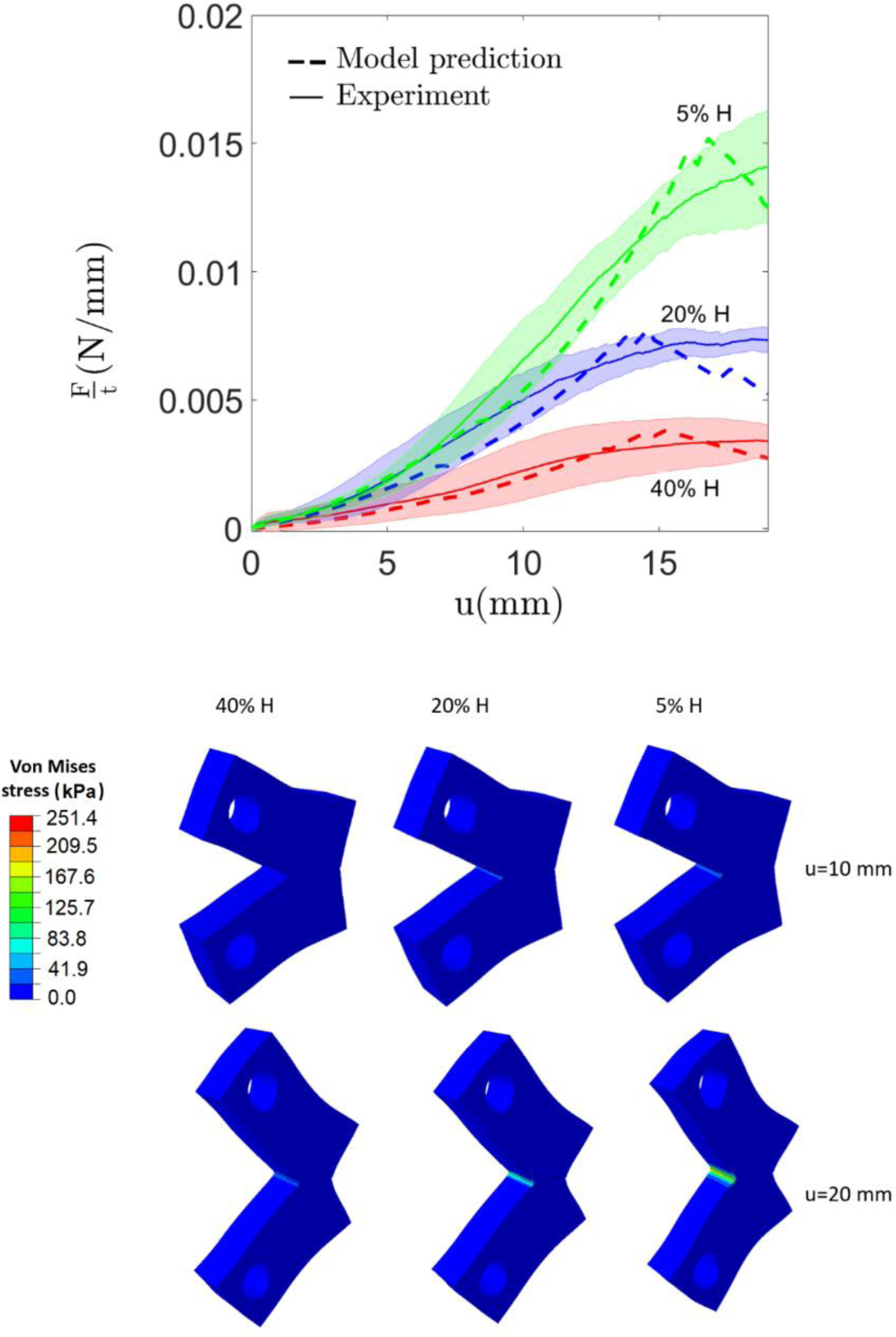
**Finite element cohesive zone model predictions of fracture initiation and propagation** for 5% H, 20% H, and 40% H platelet-contracted clot analogues. Anisotropic hyperelastic material parameters shown in Table 3 have been used.

## 4. Conclusions

This study provides the first characterization of the fracture toughness (tear resistance) of blood clot analogues. The results of which are highly relevant for assessment of clot fragmentation in thrombectomy, a major risk during endovascular treatment of AIS. A bespoke experimental test-rig and compact tension specimen fabrication has been developed to measure fracture toughness of thrombus material. Fracture tests have been performed on three physiologically relevant clot compositions: a high fibrin 5% H clot, a medium fibrin 20% H clot, a low-fibrin 40% H clot. Fracture toughness is observed to significantly increase with increasing fibrin content, i.e. RBC-rich clots are more prone to tear during loading compared to the fibrin-rich clots. This observation is consistent with the published clinical data which report high RBC-content clots in patients with multiple secondary embolism resulting from periprocedural thrombus fragmentation [20,24].

Finite element cohesive zone simulations of fracture tests are performed to further investigate the toughening mechanism associated with the clot’s fibrin network. Simulations reveal that, with increasing applied displacement, fibrin fibres behind the crack tip reoriented toward the tension direction, becoming highly aligned perpendicular to the crack direction. This significantly increases the computed J-integral at the point-of fracture initiation. Furthermore, our investigation reveals that the anisotropic hyperelastic contribution of the fibrin network must be incorporated into a computational model in order to accurately simulate experimental behaviour in both tension and compression. In contrast, an isotropic hyperelastic formulation does not capture both compressive and tensile behaviour. This insight into the material model is an advance on our previous isotropic models [27,28], which were proposed based on only considering compression experiments. Also, the CZM modelling and J-integral analysis have only been used for a limited number of studies of biological soft tissues [52,53], and have not been used for blood clots before. The developed computational model provides a suitable basis for pre-clinical assessment of the stent-retriever devices and thrombectomy procedure.

In the current study, we have investigated the influence of clot composition (fibrin and RBC) on the fracture toughness of platelet-contracted clot analogues for three compositions. There are many other compositions/contractility levels that should be considered in future studies. Moreover, the influence of recombinant tissue plasminogen activator (rtPA) on fracture toughness of clot should be considered in future experiments. Using rtPA as an early reperfusion therapy for AIS is recommended by American Heart Association and American Stroke Association (AHA/ASA) to reduce or prevent brain infarction. The experimental-computational approach that we have developed in this study could uncover the influence of rtPA on fracture toughness and fragmentation risk in vivo during subsequent MT.

## Acknowledgements

This work has been funded by a European Union Horizon 2020 Research and Innovation Program, under grant agreement No. 777072.

## Data availability

The raw/processed data required to reproduce these findings cannot be shared at this time as the data also forms part of an ongoing study.

## Appendix A: Influence of specimen thickness on the fracture toughness

The critical strain energy release rate (G_IC_) of an specimen depends on the thickness of the specimen [44]. What is referred to as the fracture toughness is the value of the critical strain energy release rate for the specimen which is thick enough to ensure the establishment of plane strain condition at the crack tip. To investigate the influence of specimen thickness on the fracture test results, we performed a series of CZM simulations for clot of different thickness. The results, as shown in Figure A1, reveal that the thickness of the tested samples (Table 2) are high enough for a valid fracture test.

**Figure A1:**
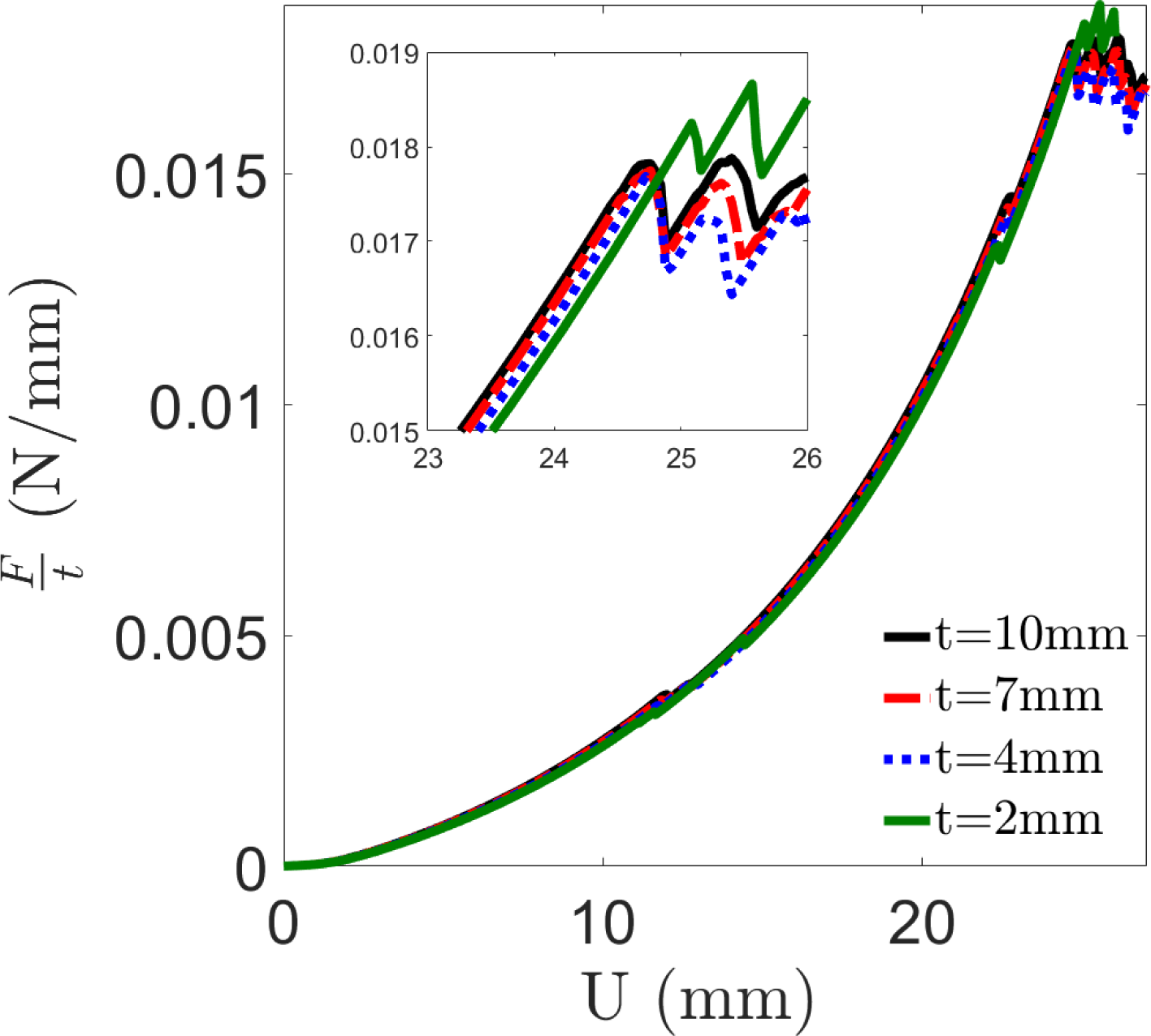
**Influence of specimen thickness on the fracture test results** for a 5% H clot analogues.

## Appendix B: Role of fibrin fibres in tension-compression asymmetry of clot

In this appendix we show that the traditional Ogden hyperelastic model with asymmetric tension-compression behaviour is not able to capture both the tension and compression test results, further highlighting the key contribution of the re-alignment of fibrin fibres in tensile loadings.

**Figure B1:**
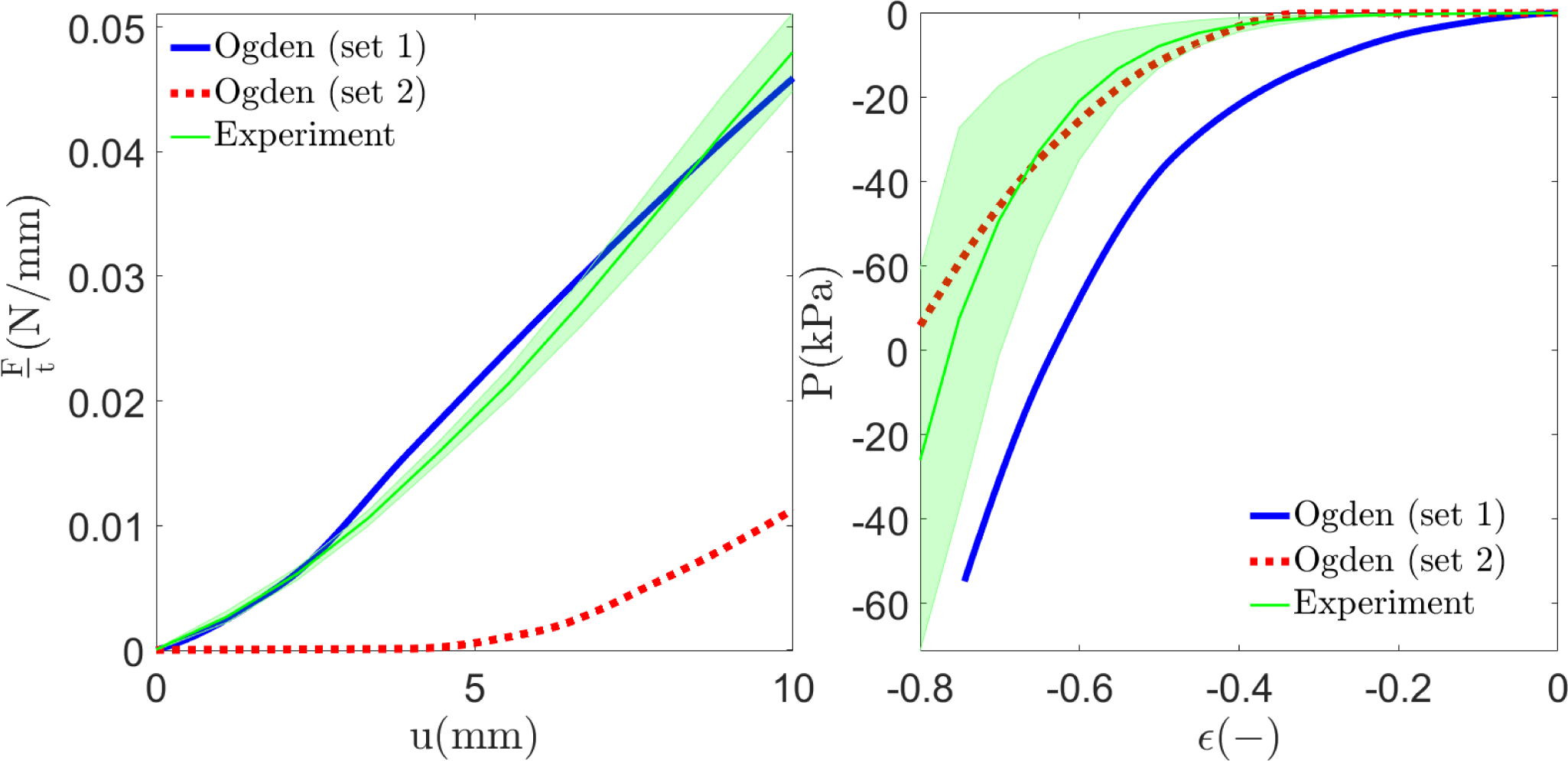
The prediction of Ogden model in unnotched compact tension specimen test (left) and unconfined compression test (right) for 5% H clot analogues. Two representative sets of parameters are visualized.

## Notes

### Competing Interest Statement

The authors have declared no competing interest.

